# Improved Temporal and Spatial Focality of Non-invasive Deep-brain Stimulation using Multipolar Single-pulse Temporal Interference with Applications in Epilepsy

**DOI:** 10.1101/2024.01.11.575301

**Authors:** Emma Acerbo, Boris Botzanowski, Damian Dellavale, Matthew A. Stern, Eric R. Cole, Claire-Anne Gutekunst, Miller L. Gantt, Melanie Steiner, Florian Missey, Antonino Cassara, Esra Neufeld, Ken Berglund, Viktor Jirsa, Robert E. Gross, Daniel L. Drane, Eric Daniel Glowacki, Andrei G. Pakhomov, Adam Williamson

## Abstract

Temporal Interference (TI) is an emerging method to non-invasively stimulate deep brain structures. This innovative technique is increasingly recognized for its potential applications in the treatment of various neurological disorders, including epilepsy, depression, and Alzheimer’s disease. However, several drawbacks to the TI method exist that we aim to improve upon. To begin, the applied electric field in the TI target is not much higher than what non-invasive transcranial alternating current stimulation (TACS) provides in the cortex. Additionally, the TI stimulation onset is dependent on the envelope of the amplitude modulated (AM) signal, where for example 1 Hz and 100 Hz envelopes have significantly different rise times to reach maximum envelope amplitude – unlike square biphasic pulses. This limitation in turn prevents classic TI, from applying bursts of pulses. Finally, the electric field intensity of TI cannot be increased or decreased at the target without dramatically altering the spatial profile of the stimulation focus. In the work presented here, we efficiently address all three of these limitations. First, we performed two-photon calcium imaging to show that individual neurons selectively respond to the TI envelope frequency, providing evidence that TI modulates neural activity with temporal specificity. This marks a significant advancement, representing the first empirical demonstration of neuronal activation at the Δf frequency within the context of TI and in an imaging modality. Subsequently, we compared the AM signals of TI with phase-shift keying (PSK) modulated signals to highlight the superior effectiveness of noninvasive pulses in contrast to the traditional TI method, particularly in inducing epileptic activity (after-discharges) in mice. We also added a multipolar configuration to create a significant increase in the electric field at the target without significantly altering the spatial profile and applied Fourier components to replicate classic biphasic bursts of square pulses - all transcranially, without the use of penetrating electrodes. These innovations aim to enhance the precision and efficacy of TI stimulation, to advance its application in neurological research and therapy.

**Key Points / Highlights:**

1. Non-invasive temporal interference stimulation modulates the activity of individual neurons at the envelope frequency.
2. A non-invasive multi-pulse TI stimulation paradigm improves both temporal and spatial focality in the deep target neural tissue when compared to traditional continuous wave (amplitude-modulated) TI stimulation.
3. Pulse TI paradigms can stimulate deep neural targets with reduced amplitude of the topical high-frequency stimulation, decreasing off-target stimulation when compared to continuous wave TI patterns. As a consequence, pulse TI stimulation reduces the risk of undesired side effects such as high-frequency conduction block in off-target tissues or cortical areas.
4. Both temporal and spatial focality of the TI stimulation pattern positively correlate with the efficacy of the stimulation to induce seizures in the mouse hippocampus.

## Introduction

Electrical stimulation as a neuromodulation method in epilepsy has been increasingly explored and included in clinical therapy and tissue exploration to investigate neural network, providing insights into the potential functions of the brain region stimulated^1,2^. As the limitations with newly developed antiseizure medications (ASMs) increases^3^, neuromodulation in the form of electrical stimulation provides an alternative treatment modality that allows selective stimulation and activation of brain structures, which cannot be targeted with medications^4,5^. The most common method of targeting deep brain structures in epilepsy is Deep Brain Stimulation (DBS) using implanted electrodes, most typically targeting the thalamus, hippocampus, or parts of the basal ganglia^6,7^. Additionally, invasive DBS is often utilized in epilepsy for investigative purposes by leveraging electrodes present for stereoelectroencephalography (SEEG), a technique consisting of implanting numerous intracerebral electrodes to assist in the precise identification of the epileptogenic zone (EZ) in patients^8,9^. Although extremely effective^10^, invasive electrode implantation can induce significant tissue damage, and the surgery itself, due to the level of invasiveness, has inherent risks^11–13^. As a result, numerous non-invasive neurostimulation techniques are currently being explored to investigate brain function and provide therapeutic stimulation^14–16^. However, the lack of focality of non-invasive stimulation methods, particularly for deep structures, remains a problem^16^.

To address this problem, a method for focal and non-invasive stimulation was proposed, temporal interference (TI) stimulation, which can non-invasively stimulate deep neural tissue, at a significant distance from the electrodes placed on the scalp. With TI, two electric fields at two different high frequencies are applied, resulting in deep, localized, and steerable temporal field envelope modulation, from constructive and destructive interferences. The modulation frequency is equal to the difference between the two applied high frequency currents, and the point in space where the modulation magnitude is maximal and stimulation is expected to occur is determined by the stimulation electrode placement and the current amplitude ratio^17^. Recently, we demonstrated the ability of TI to be pro-excitatory^18^ and pro-inhibitory^19^ in the context of an epileptic mouse model and also expanded the application of TI stimulation to peripheral nerves in animal models^20^ and the treatment of humans with obstructive sleep apnea^21^. However, several limitations of TI still exist: 1) while the higher carrier frequency permits to increase applied currents compared to transcranial alternating current stimulation (TACS) (due to reduced skin sensations and higher safety thresholds^22^), the electric field at the target is sub-threshold compared to other supra threshold technique such as Transcranial Magnetic Stimulation (TMS) – which applies much higher electric fields. In addition, 2) the stimulation onset is unclear as it is dependent on the slope of the envelope of the amplitude modulated (AM) signal, which means that stimulation envelopes of a few Hz and envelopes over 100 Hz have very different times to reach maximum envelope amplitude (proportional to the modulation period). This is unlike square biphasic pulses where stimulation is rapidly applied at maximum amplitude and the duration of the pulse can be controlled independent of the stimulation frequency, thus allowing customizable bursts. Classic TI cannot apply bursts of pulses. Finally, 3) the electric field intensity cannot be increased or decreased at the target without dramatically altering the spatial profile of the stimulation location, as classic TI only uses two current sources. Increasing the stimulation amplitude will also increase the radius of stimulated tissue and the limit is related to different factors, such as confounding skin sensation, intolerance of the subject to effects induced by the current density or charge accumulation at electrode-skin interfaces, or (at high frequencies) brain temperature increase ^23^.

In the present study, we first demonstrate the potential for TI to induce temporally-specific neural responses by measuring its effect on individual neurons using two-photon calcium imaging. We then address the above-mentioned limitations of TI by modifying the classic TI method. First, we replace the AM signals of TI with phase-shift keying (PSK) modulated signals, giving superior control over the modulation envelope slope, the pulse duration (independent of the stimulation frequency), and the ability to provide classic bursts (single pulse – spTI). We then add a multipolar configuration (multiple pairs of stimulation electrodes -mpTI) to create significant increases in the electric field at the targeted brain structure while avoiding a corresponding increase elsewhere (or even reducing off-target exposure), by essentially allowing many classic TI foci to overlap in space. By overlapping multiple TI foci in space that all produce a high exposure in the target region, but differ strongly in their collateral exposure, we effectively establish a combination that is more focal than what could be achieved with any individual electrode pair. In addition to being more focal, it is possible to significantly enhance target region stimulation intensity and/or to reduce scalp exposure and electrode current strengths, which in turn minimizes potential risks associated with tolerance issues. Finally, we apply Fourier components in the multipolar layout to replicate classic square biphasic bursts of square pulses – transforming the aggregate pulse into a square pulse, fully replicating classic DBS, but using transcranial electrodes.Utilizing a mouse model of epilepsy, we provide evidence for the effectiveness of these modified TI techniques and show that pulse-TI outperforms AM TI in inducing AfterDischarges in the context of kindling protocol.

## Methods

## 1. Animal models

Kindling and depth probe recording experiments were performed in accordance with European Council Directive EU2010/63, and French Ethics approval (Williamson, n. APAFIS#20359 -2019041816357133 v10). OF1 mice (8 and 12 weeks old) were housed in transparent cages in groups of three to five, in a temperature-controlled room (20 ± 3°C) with a 12/12h dark/light cycle. All animals had *ad libitum* access to food and water. Intravital studies were performed at Emory University with approval by the Institutional Animal Care and Use Committee in accordance with the guidelines of the National Institutes of Health’s Guide for the Care and Use of Laboratory Animals.

## 2. Intravital Two-Photon Imaging Animal Preparation

### 2.1 Viral vector

An adeno-associated viral (AAV) vector was produced in-house following a modified standard procedure^24^. In brief, human embryonic kidney 293FT cells were transfected with three plasmids (helper, AAV2/9 rep/cap, and pAAV.Syn.Flex.NES-jRGECO1a.WPRE.SV40 (Addgene, Plasmid #100853)) using the calcium-phosphate method. Recombinant AAV was harvested, purified through ultracentrifugation, and concentrated in Dulbecco’s phosphate-buffered saline supplemented with 0.001% (v/v) Pluronic F-68. It was aliquoted and stored at -80°C until surgery. The titer was determined to be 1×10^14^ viral genomes/mL via quantitative PCR.

### 2.2 Cranial Window Implantation

Stereotactic surgery was performed on an adult (≥ 90 days old) Vglut1-IRES2-Cre (Slc17a7-IRES2-Cre, Jackson Laboratory, Stock No. 023527) mouse to prepare them for chronic intravital two-photon calcium imaging of cortical pyramidal cells. The protocol for this surgery was adapted from standard protocols for concentric cranial window implantation^25,26^. A 3 mm craniotomy was performed over the anterior left hemisphere to visualize the primary motor cortex. Stereotactic injections of 300nL of adeno-associated viral (AAV) encoding the red genetically encoded calcium indicator (GECI) jRGECO1a^27^ were performed through a pulled glass capillary (Nanoject 3.0, Drummond) at a rate of 2nL/s at depths of 300μm and 600μm from the pial surface (0.30 mm anterior and 1.75 mm lateral to Bregma)^28,29^. Next, a concentric window (inner diameter 3 mm, outer diameter 5 mm; D263 #1 thickness coverglass, Warner Instruments) was placed to plug the craniotomy and affixed to the skull using dental acrylic (C&B Metabond, Parkell). A stainless steel headplate (Neurotar) was then positioned around the window and attached to the skull using dental acrylic. Burr hole craniotomies (0.5mm) were then performed bilaterally anterior and posterior to the headplate for the placement of stainless -steel skull screws with pre-soldered gold pin attachments (E363/96/1.6/SPC, P1Technologies) through which TI stimulation would be delivered. These screws were affixed to the skull using dental acrylic and the skin was closed to this headstage using tissue adhesive (Vetbond, 3M).

### 2.3 Intravital Two-Photon Imaging

Imaging was performed under general anesthesia (1.5% isoflurane balanced in oxygen, flow rate: 1 L/min) one month following surgery to allow for adequate GECI expression. The mouse was head-fixated under the objective with their snout positioned next to a nose cone for gas anesthetic delivery and a heating pad was placed under the mouse to maintain the body temperature during the imaging sessions. Resonant galvanometer scanning (30 Hz, 512 × 512 pixels) calcium imaging was performed through a long working distance 16x objective (water-immersion, N.A. 0.80, Nikon) using a two-photon microscope (HyperScope, Scientifica) equipped with a pulsed tunable infrared laser system (InSight X3, Spectra-Physics) and ScanImage (Vidrio Technologies) controller software. jRGECO1a fluorescence was excited using a 1030 nm wavelength light and fluorescence emission was separated by a dichroic with band pass filter (565LP, ET620/60m-2p Chroma) and collected using a multi-alkali red-shifted PMT. Imaging sessions lasted for 45 s and consisted of 5-10 s of baseline fluorescence imaging followed by 30 s of TI stimulation and then a 5-10 s washout. Subsequent recordings sessions were separated by at least 1 min. TI was delivered as indicated in following sections.

## 3. Simulation of the Electric Fields

### 3.1 Exposure Quantities of Interest

Given two fields at a location x, we shall name the larger one (by absolute magnitude) E^→^_1_ and the other one E^→^_2_. During most of the exposure time, they are in antiphase, producing a total field E^→^_A_ = E^→^_1_ ™ E^→^_2_. During the pulse, they are in phase, resulting in E^→^_I_ = E^→^_1_ + E^→^_2_. It is assumed that |E^→^_I_| ≥ |E^→^_A_| (i.e., the pulses are peaks, rather than dips). The below argumentation can easily be adapted for the other (less relevant) case. The exposure difference between in- and anti-phase is then E^→^_I_ - E^→^_A_ = 2E^→^_2_. A neuron with axis alignment to E^→^_2_ would experience a change of 2 |E^→^_2_| in projected field exposure, i.e., twice the smaller field amplitude.

Then, the highest momentaneous high frequency field magnitude for pulsed TI is |E^→^_1_|. Classic TI goes through all phase relationships between the two fields and also reaches its biggest magnitude – namely |E^→^_1_| – when they are in phase. However, the smallest magnitude is not always |E^→^_A_|. Instead, when the projection of E^→^_A_ along E^→^_2_ points in the opposite direction of E^→^_2_, the minimum absolute field magnitude is reached for an intermediate phase between 0° and 180°. In that case, the biggest modulation amplitude is not twice the smaller field magnitude anymore.

In that case, it can be shown, that the neuron orientation which experiences the maximal modulation is perpendicular to E^→^_A_ and co-planar to the plane spanned by E^→^_1_ and E^→^_2_. It is the direction along with the projection along that neuron orientation becomes zero, when the fields are in antiphase. The formula given in Grossman et al., 2017 computes these magnitudes for the two cases. As long as the quantity of interest is the modulation amplitude along the orientation where that amplitude is maximal, the same formula also applies for pulsed TI.

However, there are alternative quantities that could be evaluated and that are not identical for pulsed and classic TI. For example, the modulation magnitude of | E→*(t)* |. For pulsed TI, it becomes||E^→^_I_| - |E^→^_A_||, for classic TI it is |E^→^_I_| - |E^→^_I_ - E^→^_2_ ((E^→^_I_. E^→^_2_)/ (|E^→^_I_| |E^→^_2_||). Which quantity is more suited (in terms of correlating with the resulting impact on neural dynamics) is debatable and depends on the yet unclear interaction mechanism(s).

### 3.2 Simulation of Electric Fields in Mouse Model

The simulation of electric fields in a mouse model was conducted using the online accessible Temporal Interference Planning Tool (TIP) V2.0, an online tool developed by IT’IS Foundation. This tool, powered by o^2^S^2^PARC technology^30^, allowed to define and optimize targeted neurostimulation protocols in a three-step process. First, the target brain regions and constraints on potential electrode placement are defined. Next, an optimized stimulation protocol is identified through electromagnetic simulations involving detailed anatomical head models, automated multi-goal optimization (maximize target exposure strength, target coverage, and avoidance of collateral stimulation), and interactive refinement (i.e., adaptation of the weights of the three goal functions, resulting in the immediate identification and visualization of a new strategy from the previously determined Pareto front). A detailed report with both quantitative and visual information was then generated. The exposure analysis of the selected parameters was performed using the Sim4Life web platform. The TIP V2.0 tool offers a comprehensive library of highly detailed anatomical head models, including a mouse model for studies involving rodents. It also provides options for multichannel TI and phase-modulation TI, enhancing flexibility in stimulation parameters. Here, the coordinates for the stimulation electrodes were previously published^18^, and the closest pairs recommended in the simulation were selected to replicate the electric fields using sim4life (here : CP1 to CP2 and CP3 to CP4).

## 4. Surgical procedure for TI stimulation (awake and anesthetized)

### 4.1 Surgical implantation for awake recordings

Mice (n=9) underwent a surgical procedure to implant minimally invasive screws to perform TI stimulation. First, mice were anesthetized via an intraperitoneal injection of a mix of Ketamine (50mg/Kg) and Xylazine (20mg/Kg) before being placed in a stereotaxic frame. After a midline incision, a verification that the lambda and bregma were on a horizontal plane was done before getting bregma coordinates. Craniotomies for the placement of minimally invasive stimulating electrodes (2 pairs of stainless steel mini-screws, Component Supply, Miniature Stainless Steel Self-Tapping Screws: TX00-2FH) were performed at [AP: -1.94, ML: +0.5; -0.5; -3.9; -4.3] as described previously^18^ and a depth probe was introduced into the hippocampus (Implantable twisted-pair platinum electrodes from PlasticsOne; wire length = 5mm, individual wire diameter = 125µm) at [AP: -2.7, ML: +2.04, DV: 1.30] using a 20-degree angle. Cortical electrodes were screwed into the skull with minimal penetration into brain tissue. A reference for the depth electrode was placed in the cerebellum. Then, to fix all the implanted electrodes, dental cement was applied (Phymep, SuperBond).

### 4.2 Surgical implantation for anesthetized recordings

One mouse was deeply anesthetized (intraperitoneal injection of ketamine (50mg/kg) and xylazine (20mg/kg)) and implanted with a multiple recording sites (32) electrode (NeuroNexus A1×32 Edge 5mm-177, length: 32mm, contact-spacing:100µm) in the hippocampus [AP: -2.7, ML: +2.04, DV: -3.2]. We also performed multiple craniotomies to implant 8 pairs of screws to surround the depth probe and create a focus of stimulation at the center. This experiment was an acute experiment due to the high number of electrodes placed in a single mouse.

## 5. Recordings and Stimulation

### 5.1 Electrical stimulation

Standard TI patterns were implemented by simultaneous excitation of the tissue with two sinusoidal signals of frequencies 1000 Hz and 1050 Hz. The linear superposition of these two signals in the neural tissue produces a continuous envelope at 50 Hz with a sinusoidal shape. To improve the spatial focality of the interference pattern on the target neural tissue, the TI paradigm was also implemented using a multipolar TI (mTI) montage in which multiple pairs of interfering sinusoidal signals are applied simultaneously to produce the AM envelope at the deep target.

In addition, two novel forms of TI were developed to improve both temporal and spatial focality relative to the standard TI stimulation. The first is single-pulse TI (spTI) which produces a train of sinusoidal pulses generated at the tissue target via the interference of two signals: a) a pure sinusoidal tone at 1000 Hz and b) a sinusoidal signal of the same frequency (1000 Hz) with phase inversions at a rate equal to the desired stimulation frequency. As a result, the linear superposition of the unmodulated carrier (a) and the PSK modulated signal (b) produces an interference deep in the neural tissue, resulting in a train of pulsed field changes at the phase inversion rate. In the case of mpTI, multiple spTI-modulated signals are applied simultaneously to produce a train of sinusoidal pulses at the deep target tissue. Finally, a stimulation paradigm introduced in this work allows for generating a train of flexibly shaped pulses deep in the neural tissue. This novel approach is based on the Fourier series representation of the desired waveform shape and can be implemented using both continuous (e.g. Phase Modulation) and discrete (e.g. PSK) modulation paradigms.

To perform stimulation, waveform generators (EDU33210A Keysight, USA) were driving independent current sources (DS5s Digitimer, UK), which were injected via the implanted cortical electrodes.

Kindling Protocol: After a few days of recovery, mice were connected to the recording and stimulation setup. Stimulations were applied on the cortex by means of two DS5 current sources driven by one function generator (with two independent channels). Here, the aim was to determine the after-discharge threshold (AD threshold) for spTI vs. regular TI using sine waves. Mice were connected and placed back in their cages. The frequency (Δf) was 50Hz in both conditions. The amplitude of stimulation was increased in steps of 50µA until the triggering of an epileptic seizure as described in Missey et al. 2021^18^ and Acerbo et al. 2022^19^.

Depth Probe recordings using NeuroNexus Probe: We also wanted to assess the focality of the different techniques of TI stimulation. We focused on the maximum amplitude of exposure at the end of the NeuroNexus probe to see if we could determine the amplitude of the stimulation at the focus. We recorded the artifact of stimulation via the 32 electrodes on the probe connected to the recording and stimulating device. Since we were working on an anesthetized mouse, we could add several pairs of stimulation electrodes. We thus also analyzed the combination of 8 pairs of stimulation electrodes creating mTI. Further explanations regarding this improvement of the TI method can be found in Botzanowski et al. 2023^31^.

### 5.2 Recordings

All electrophysiological recordings were performed with a stimulation/recording controller (IntanTech, Intan 128ch Stimulation/Recording Controller) with a sampling rate of 30kHz. To perform stimulation, waveform generators (EDU33210A Keysight, USA) were driving independent current sources (DS5s Digitimer, UK), which were stimulating via the implanted cortical electrodes.

## 6. Data and Statistical Analysis

All the recordings were initially saved in the .rhs format and later converted into MATLAB files for data processing and statistical analyses.

### 6.1 Two Photon Imaging

Motion registration, region of interest (ROI) detection, and transient extraction of soma and surrounding neuropil were performed using the Suite2P software package^32^ with integrated Cellpose^33^. All calcium transient data was then processed and analyzed in MATLAB. The raw fluorescence traces were background subtracted and 70% of the surrounding neuropil signal was subtracted from the somatic signal to remove the out-of-plane signal from the surrounding neuropil^32^. These traces were then normalized as dF/F_0_. Additionally, mean-field fluorescence traces were extracted using FIJI^34^, the background was subtracted, and the signal was normalized as dF/F_0_. For all recordings, F_0_ corresponded to the first 5 seconds of baseline recording before stimulation.

Power spectral densities were computed within each cell using Welch’s power spectral density estimate and normalized to the total power within cell. While all analysis was performed on unfiltered fluorescence data, the traces presented are double reverse filtered using an adaptive filtering routine (Butterworth lowpass filter, order 3 to 5) that preserves the phase of the signal (mean field: 2 Hz; individual cell: 1 Hz). Analysis was performed on the top half of cells by relative power in the frequency range of interest (<1.5Hz). Power distributions across cells were tested for normality using the Kolmogorov-Smirnov test. As these data were found not to be normally distributed, non-parametric statistics were used, namely the Wilcoxon rank sum.

### 6.2 In vivo recordings

The statistical tests for the behavioral experiments were done using R Studio®. For each experiment, a test on the data distribution was performed. In this case, the distribution of the AD threshold for each mouse was normal. Thus, t-tests (parametric, paired) were performed to determine if a groups’ distribution data was different.

To assess the amplitude of the envelope in the case of the depth probe multi-site recording experiment, we analyzed the signal in MATLAB over a 10-second interval. The envelope of the signal was obtained using the envelope function, and 20ms windows were created to calculate the (max-min) envelope values. Subsequently, the median value and standard deviation were calculated across the windows. In order to test the distribution of TI versus spTI, we performed an Anderson -Darling test to evaluate, whether they follow normal distributions. Since only spTI exhibited a normal distribution, we used a rank-sum test to compare the two groups. For the Fourier stimulation, amplifier saturation occured for 3 contacts (N°4-5-6) necessitating data point interpolation to preserve the continuity and overall shape of the data while mitigating the impact of outliers. We employed cubic spline interpolation with the ‘spline’ option in MATLAB to estimate the missing values. The interpolation procedure aimed to maintain the general trend of the data while filling in the gaps caused by missing values. Finally, to test for the time focality, we used the findpeaks function to determine the width of each peaks. For TI and mTI signals, we used their envelope to calculate the width.

## Results

### Neuronal Responsiveness to Temporal Interference

To investigate the impact of TI on individual cell responses, we conducted intravital two-photon calcium imaging through chronically implanted cranial windows in an anesthetized mouse. Imaging was targeted to pyramidal cells in cortical layer 2/3 using a red GECI expressed in VGLUT positive cells (Figure 1A-B). TI was applied using two dipoles generated from stainless steel skull screws positioned outside the cranial window, with a carrier frequency of 2 kHz. Envelopes below 1 Hz were selected to accommodate the temporal resolution limitations imposed by GECI kinetics. Notably, we observed responses throughout the cortical field, including soma and neuropil, that were entrained to the envelope frequencies of stimulation (Figure 1C).

**Figure 1:**
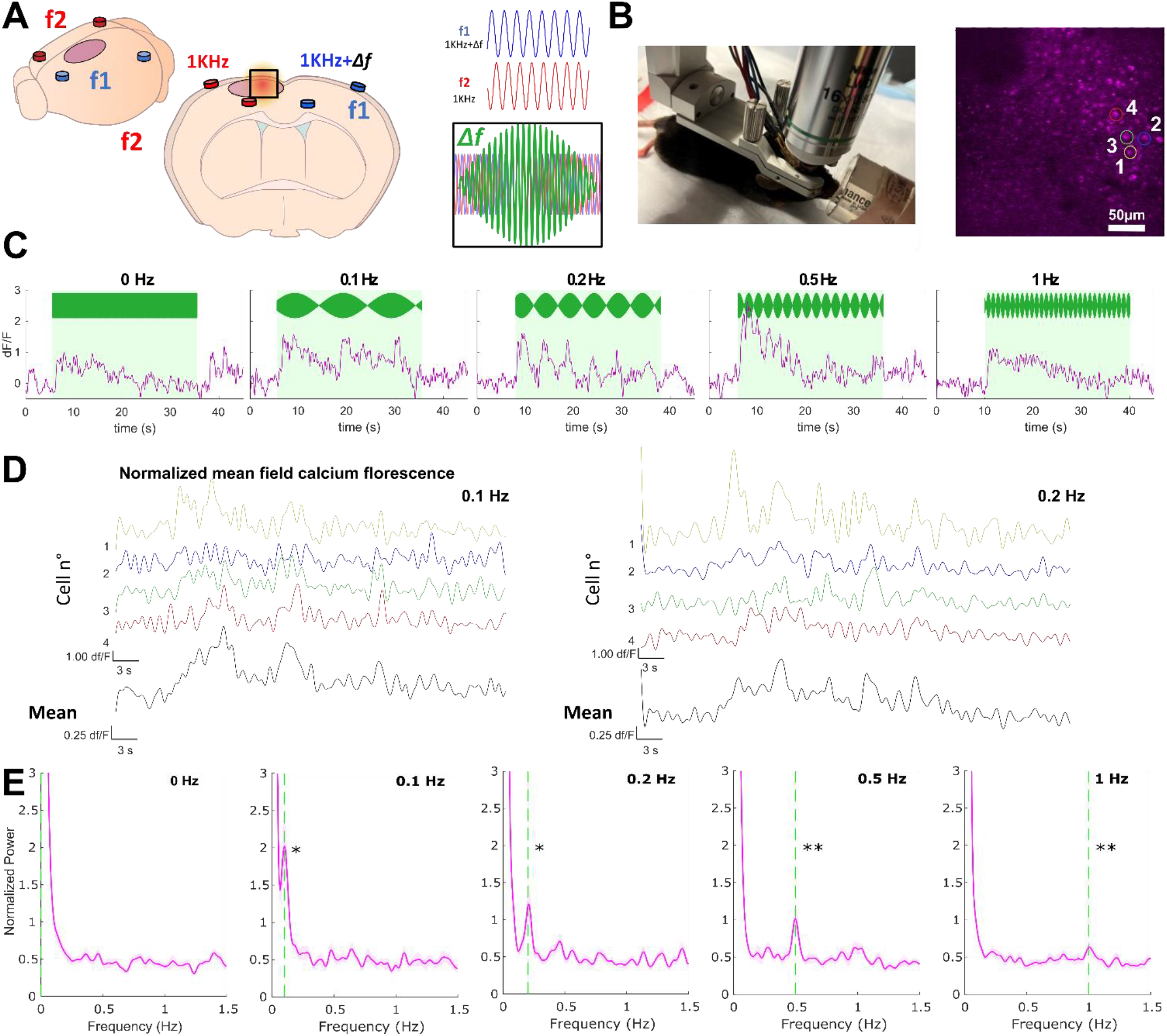
Temporal Interference Modulates Cortical Neurons at Envelope Frequency. A) Principle of temporal interference (TI) stimulation. B) Image of the experimental setup for in vivo two-photon calcium imaging in an anesthetized mouse during TI stimulation. The headstage montage includes screw pairs (color-coded) for generating TI stimulation (2 kHz sine wave carrier frequency, 0-1 Hz envelope frequencies, 400 µA amplitude). The image shows the headplate and cranial window, with the average field of view of layer 2/3 pyramidal cells (jRGECO1a::VGLUT1) highlighted. Scale bar (white) represents 50 µm. C) Normalized mean field calcium fluorescence for different TI stimulation envelopes. The green box indicates the stimulation period with the overlaid stimulation pattern. D) Normalized calcium fluorescence traces in selected individual cells (labeled by name and color in panel B), with the average trace of all extracted cells (N=48-72 cells) shown below in black. Scale bars:1.00 dF/F by 3 s. E) Average power spectral densities across recruited cells (µ±sM) isolated for each stimulation envelope, with the dashed line indicating the stimulation envelope frequency. Wilcoxon rank sum analysis compares the power at the stimulation envelope frequency to the power in the carrier frequency alone condition (0 Hz). *p<0.05, **p<0.01.

At the individual cellular level, we further observed that the somatic calcium fluorescence of single cells was also modulated at the envelope frequencies (Figure 1D-E). These experiments collectively confirm that TI can induce and modulate neuronal activity at envelope frequencies at the individual cell level. However, an essential finding of our study highlights the ambiguity surrounding the precise timing at which the envelope of stimulation triggers a detectable event. While we provide evidence that neurons are responsive to the frequency of the envelope, determining the specific phase at which neurons respond within the envelope remains challenging.

To address this limitation, we have developed pulse TI (spTI), a technique that offers enhanced temporal precision. This approach bears resemblance to pulse protocols employed in electrical stimulation mapping for epilepsy or tumor cases involving SEEG electrodes^35^. Notably, pulse width can be modified by adjusting the carrier frequency, enabling us to maintain the same envelope frequency while having different pulse widths (Figure 2). The development of pulse TI provides a potential solution to enhance temporal precision in future studies.

**Figure 2.**
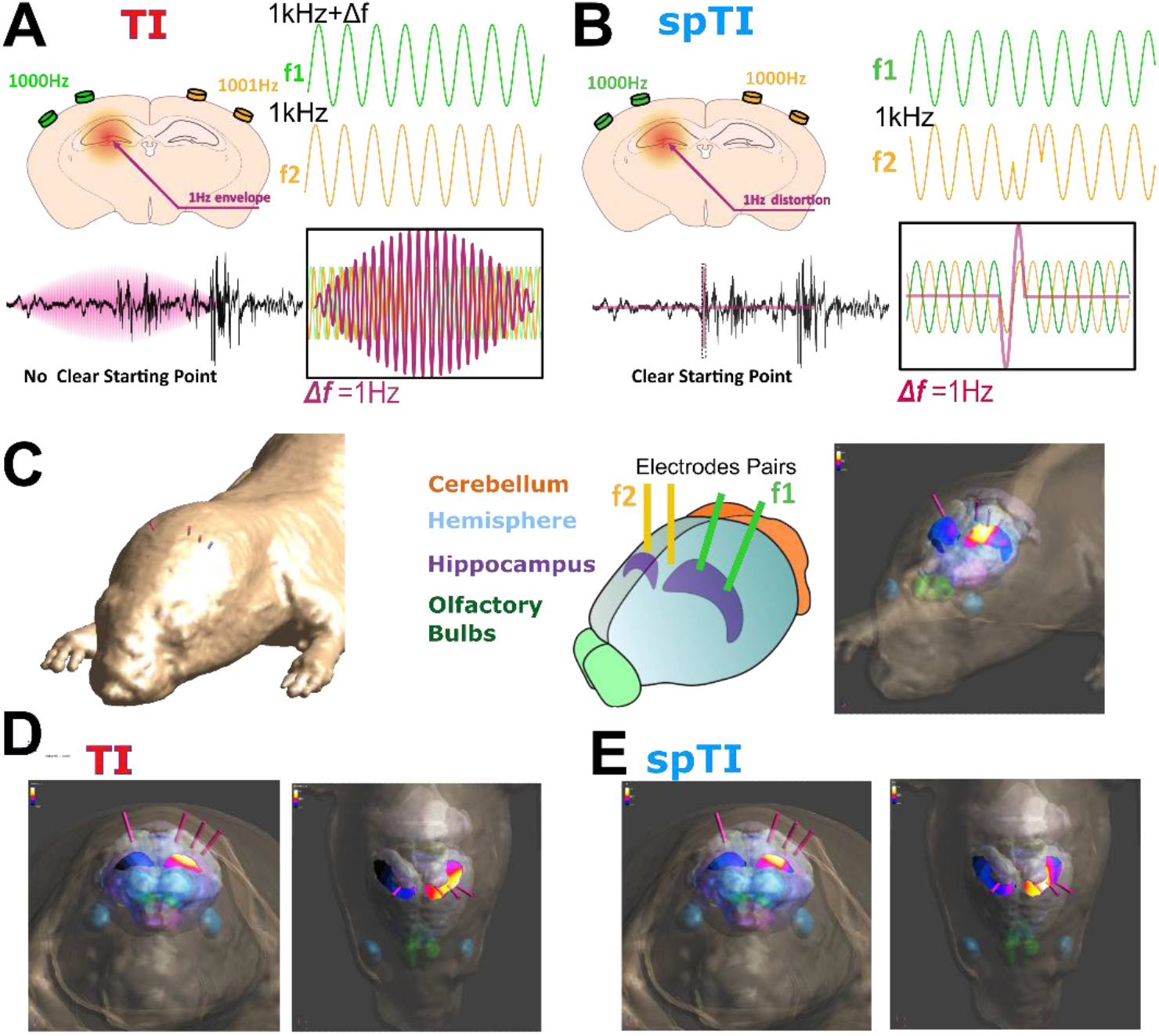
In Silico Model of Temporal Interference (TI) and Spatial-Phase Temporal Interference (spTI) Stimulation in the Hippocampus. A) Illustration of Temporal Interference (TI) in the hippocampus via two pairs of electrodes running at 1 kHz and 1,001 Hz with a stimulation frequency of 1 Hz as an example. The low-frequency amplitude modulation at 1 Hz obscures the exact timing of TI-induced neural activity. To address this, we introduced spTI, which provides a more precise temporal definition by creating a non-invasive pulse. B) Illustration of spatial-phase Temporal Interference (spTI) in the hippocampus via two pairs of electrodes running at 1 kHz and 1 Hz, with a 1 Hz phase inversion as an example. In contrast to TI, the pulsed nature of spTI provides a clearer starting point for the stimulation. C) In silico model of the mouse brain featuring various anatomical regions. Electrode pairs CP1 -CP2 and CP3-CP4 (sim4life) were chosen for stimulating the hippocampus. The right panel depicts the simulation of TI exposure in the hippocampus. D) and E) are simulations for TI with amplitude modulation (AM) and spTI, respectively. Notably, with the same current amplitude (and a 1:1 current ratio between the pairs), spTI exhibits greater focus, primarily targeting the central portion of the hippocampus, while TI stimulates the entire hippocampus.

### Introducing a New Paradigm of Stimulation: Non-Invasive Single Pulse Stimulation (spTI)

TI stimulation, utilizing AM, serves as the initial focus of this study. In our illustrations (Figures 2A and B), we first explore TI’s potential, highlighting its application in the hippocampus. In Figure A, we depict TI in the hippocampus applied via two pairs of electrodes operating at 1 kHz and 1,001 Hz, with a stimulation frequency of 1 Hz. However, one notable limitation of this technique becomes apparent: the stimulation onset is not precisely defined due to the continuous, oscillating nature of the 1 Hz amplitude modulation. The ambiguous stimulation initiation of classic TI presents a challenge in achieving precise temporal control. Responding to the need for more precise temporal stimulation, we introduce a novel approach called spTI stimulation. Figure 2B illustrates the spTI stimulation artefact in the hippocampus using two pairs of electrodes, with frequencies of 1 kHz and 1 Hz, incorporating a 1 Hz phase inversion. The introduction of this phase inversion significantly enhances the temporal precision of the stimulation, establishing a sharp reference and phase relationship (Figure supplementary 1). The *in silico* mouse brain model (Figure 2C) allows the comparaison of both TI and spTI techniques. Subsequent simulations in Figures 2D and 2E emphasize a slightly more spatial advantage for spTI compared to TI, (with equivalent current amplitudes and a 1:1 current ratio between electrode pairs). spTI effectively targets the central region of the hippocampus while TI results in more diffuse stimulation across the entire hippocampus. These results highlight the potential superiority of spTI in terms of spatial and temporal control in deep brain stimulation, particularly within the hippocampus.

### Behavioral Investigations of spTI: Comparing its Efficacy with Regular TI

Here, we compared the effectiveness of spTI and TI in inducing afterdischarges (AD) in mice, first step of a kindling protocol. The experimental configuration for the behavioral experiment necessitated the insertion of minimally invasive screws (operating at frequencies f1 and f2) into the skull. Additionally, a depth probe was implanted into the hippocampus to facilitate data collection (Figure 3A). In the classic TI stimulation (referred to as TI), we employed the following frequencies: f1=1kHz, f2=1.05kHz, for Δf=50Hz, with both frequencies recorded in the target region (hippocampus). Power Spectrum Analysis (PSD) show the two stimulation frequencies without any peak at 50Hz. This absence is due to 50Hz not truly existing as a component. (Figure 3B) However, as demonstrated in the above-presented results, we can emphasize that neurons are responding at this frequency. In the context of spTI, the frequencies utilized are f1&f2=1kHz, and one peak at 1kHz is present in the corresponding PSD. However, in this case, the phase flip is an event occurring at 50Hz and does exist, making it visible on the PSD (Figure 3C). Each mouse underwent both spTI and TI stimulations. This design allowed us to eliminate inter-subject variability and focus on AD threshold analysis. Notably, our results showed that spTI required significantly less current to induce AD compared to TI. The hippocampal AD threshold was reached with approximately 400 μA less (∼30%) current than in the case of spTI (Figure 3D). These findings underscore the efficacy of spTI in achieving similar outcomes to TI but with a considerably lower current requirement. Furthermore, spTI enhances temporal precision during stimulation, making it a valuable non-invasive neuromodulation technique for replicating clinical protocols and precisely time the onset of desired outcomes following stimulation.

**Figure 3.**
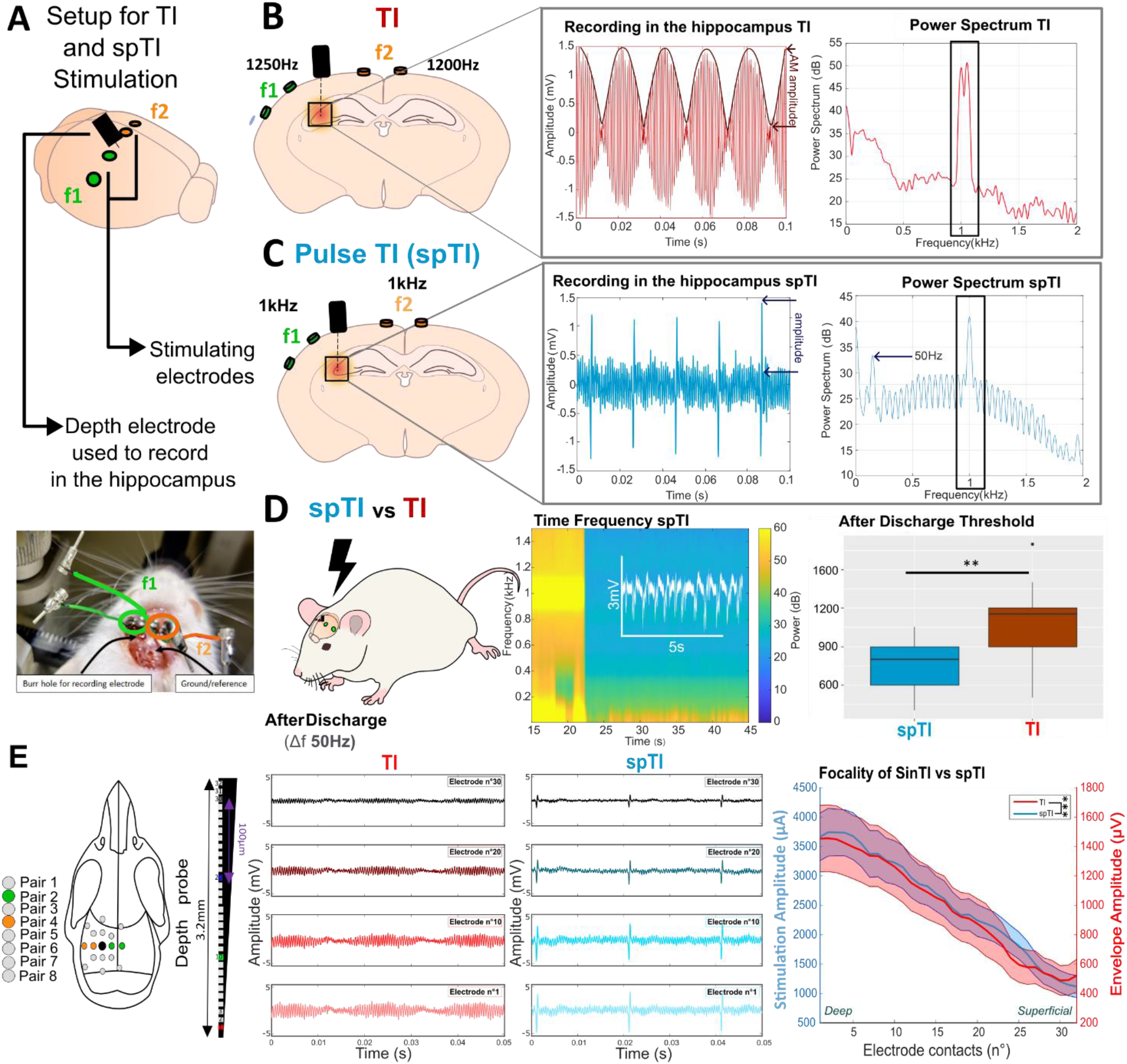
Comparison of spTI and Regular Sine Wave TI in After discharge Threshold (AD) Evaluation. A) Illustrations of experimental setup for the TI and spTI is displayed. Minimally invasive screws were placed in the skull, and a depth probe was inserted into the hippocampus for data collection. B) Temporal Interference (TI) with Amplitude Modulation (AM) recorded in the hippocampus (Δf=50) is generated by combining two high-frequency components (f1:1250Hz and f2:1200Hz). These two frequencies are detectable on the Power spectral Density (PSD) of the recorded signal from the depth probe in the hippocampus. C) spTI stimulation is applied using two pairs of electrodes on the cortex, using the same setup as in A. In the case of spTI, the same frequencies are utilized (1 kHz), but the signal’s phase is flipped every 20 ms to achieve constructive interference at 50 Hz. On the raw recording from the hippocampus a pulse at 50Hz is evident. The PSD reveals a peak at 1 kHz and another one at 50Hz. D) The threshold for Afterdischarges (AD) induced by stimulation in the hippocampus of behaving mice (n=9) at Δf=50 is analyzed. On the time-frequency plot, an AD is elicited after stimulation at 600 µA, and the threshold is determined. Time-frequency plots and raw recordings from the hippocampus are employed to identify the threshold. More current is required to evoke an AD using TI compared to spTI (p-value**=0.005). Associated behavioral videos can be found in the supplements. E) Surgical procedures were conducted on an anesthetized mouse to implant multiple cortical electrodes. Pair 2 (green) and pair 3 (orange) are used to focus TI in the hippocampus, with a recording electrode implanted in the center (black). The mouse receives stimulation using TI and spTI. Recordings are made from contact 30 (close to the cortex) to contact 1 (deepest), providing insight into the spatial distribution of the stimulation over 3.2 mm. A significant difference is observed between TI and spTI artefact stimulation amplitude (p-value***=6.2813e-10), with the shaded area representing the standard deviation.

### Comparison of regular Sine Waves TI and spTI

Having demonstrated that spTI exerts a stronger effect on the onset of epileptic seizures, we sought to explore potential additional advantages of this technique, such as improved focal stimulation. To investigate this, we performed recording on an anesthetized mouse using a depth electrode with 32 contacts spaced at 100µm (3.2 mm total electrode length). Placing it between pairs 2 and 4 of the stimulation electrodes (as depicted in green and orange, respectively), we aimed to replicate the experimental setup employed in the previous study^18^. Subsequently, we applied identical stimulation conditions (TI and spTI) to these two pairs. With the depth electrode in place, we were able to record the amplitude of the TI and spTI exposure across different recording sites.

By analyzing the recorded data, we observed the amplitude for both conditions along the contacts, ranging from the deepest (contact n°1) to the closest to the cortex (contact n°30). In the case of TI, the highest amplitude was observed at the deepest contacts, although there was minimal variation between contacts 1 and 10. In the case of spTI, the highest signal amplitude was also observed at the deepest contacts, but a visible difference in amplitude between contacts 1 and 10 was noted (Figure 3E).

To examine the stimulation profile with depth, we calculated the amplitude of stimulation for both conditions. The distribution of envelope amplitudes showed distinct patterns for TI and spTI. With TI, we observed a threefold decrease in AM amplitude within a 3.2mm range, going from 1500µA to 500µA. In contrast, spTI demonstrated a decrease in signal amplitude from 3800µA to 1000µA. This difference in amplitude offers more flexibility when using spTI, as lower applied amplitudes permit a reduction of high-frequency carrier amplitude throughout the entire brain. Significant differences in amplitude between TI and spTI were observed within various contacts, as indicated by a p-value*** of 6.2813e-10.

In conclusion, our findings highlight the potential advantages of spTI over TI, including improved focal stimulation and the ability to achieve comparable effects with lower amplitudes.

### Comparison of the different types of multipair TI

Advanced TI paradigms were developed to enhance spatial resolution of TI stimulation, as described in a previous work^36^. Consistent with expectations, the mTI paradigm, utilizing multiple pairs of electrodes, resulted in higher exposure amplitudes compared to spTI (Figure 4A). Moreover, this multipolar design was explored for two variations of spTI: mpTI and Fourier TI.

**Figure 4:**
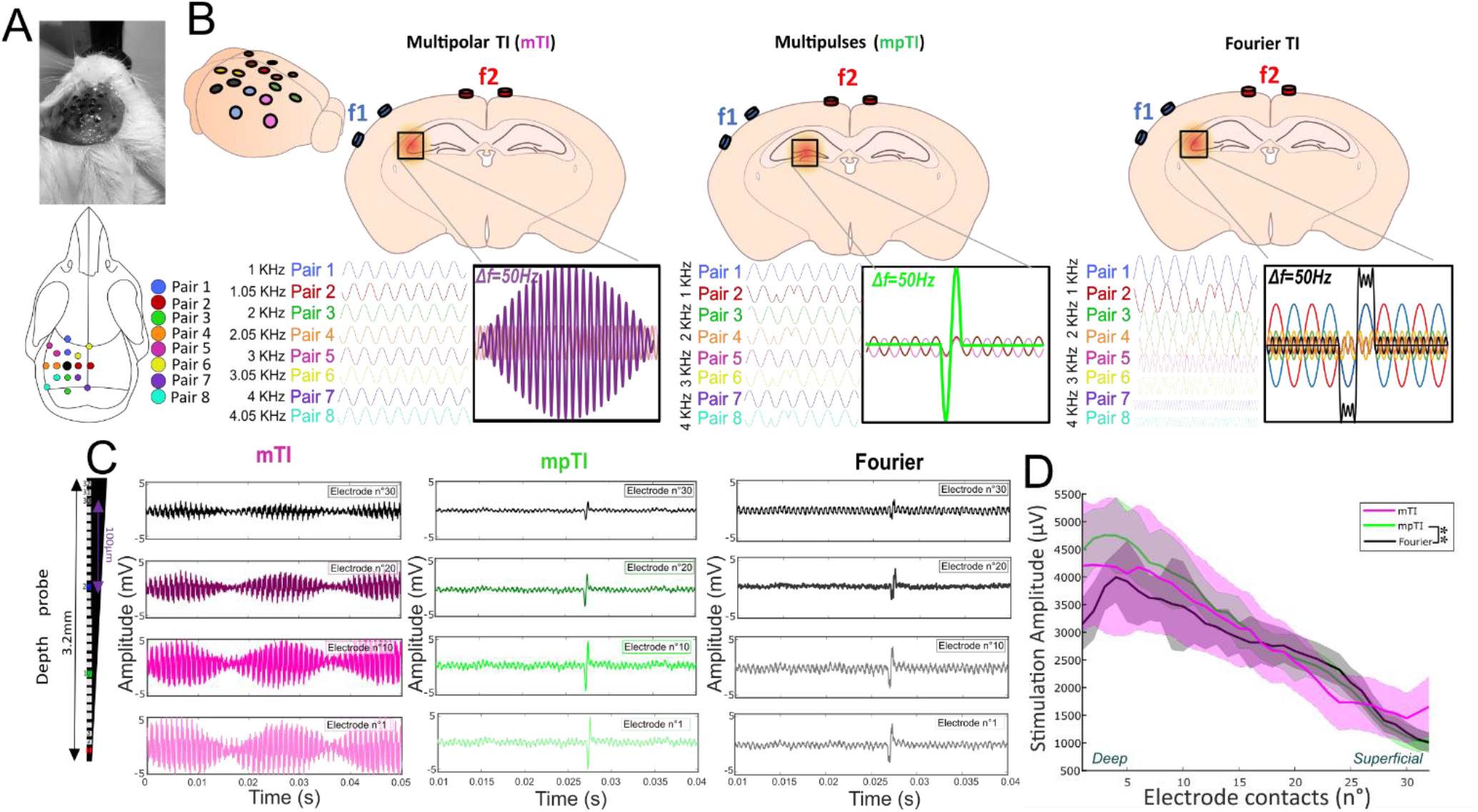
Comparison of the focus and temporality of different types of multipair TI. A) An anesthetized mouse underwent surgery with multiple cortical electrodes implanted. All the electrode pairs were utilized to generate mTI (multipolar temporal interference), Fourier, or mpTI (multipolar pulses TI). B) Illustration of the different types of stimulation using eight sine waves at different frequencies for mTI, four sine waves and their PSK waves for mpTI, and sine and PSK waves at different amplitudes for Fourier. C) The recording electrode with multiple contacts (32) allowed for the assessment of the spatial distribution of stimulation across a depth of 3.2 mm. The graph displays the stimulation patterns for mTI (pink), mpTI (green), and Fourier (black), from the closest to the cortex (electrode 30) to the deepest (electrode 1). D) No significant difference was observed in the amplitude of the stimulation artefact between Fourier and mTI (p-value = 0.48) and mTI vs mpTI (p-value=0.63). However, a significant difference was found between mpTI and Fourier (p-value < 0.001).

By employing mpTI, we overlapped four pulses generated from four PSK sinusoidal waves at depth. This composite pulse exhibited comparable characteristics to the initial spTI but with a significantly broader amplitude (while maintaining the same stimulating amplitude). Another method for achieving pulse stimulation via TI is through Fourier TI. In this approach, we overlapped four different sine waves with varying weights to generate a square-like pulse similar to those achievable with a standard implanted DBS electrode at depth (Figure 4B). Comparing these deeply evoked Fourier TI and mpTI with the mTI paradigm, we observed a substantial reduction in temporal width (while maintaining the same amplitude stimulation) (Figure supp 1, Figure 4C) as we see by comparing spTI and TI. Furthermore, we aimed to assess the spatial focality of these three multipolar TI designs. Consistent with our previous study^36^, the multipolar TI approach significantly improved the spatial focality of the stimulation by incorporating different pairs of TI with at least a 1kHz difference between them. Notably, we observed no significant difference regarding the amplitude of the stimulation between mTI and mpTI (p-value = 0.6336), indicating similar outcomes in terms of spatial focality. However, the comparison between mpTI and the Fourier revealed a significant difference (p-value = 0.0016), highlighting the advantage of the mpTI paradigm (Figure 4D). Thus, we demonstrated that mpTI exhibited a larger stimulation amplitude for the same injected current, as confirmed by the corresponding p-values. Overall, our findings emphasize the versatility of the multipulse paradigm, which incorporates spTI to generate pulses while integrating the mTI approach for precise and focal stimulation, offering promising prospects for targeted and accurate neuronal stimulation.

## Discussion

In this study, we introduce an innovative approach called spTI stimulation, designed to address the limitations inherent to traditional TI stimulation and unlock its full potential for clinical research. We first observed that individual neurons exhibit modulation at the frequency of the AM (Δf) generated through TI. This represents the first empirical observation of neuronal activation at the Δf frequency using an imaging modality that is free for stimulation artifact. Although, when scrutinizing the Power Spectral Density (PSD) of the TI waveform, no Δf frequency is discernible, in stark contrast to spTI. The significance of this observation lies in the inherent challenges posed by the characteristic frequencies of stimulation. In theory, it is possible to eliminate the stimulation artifact by applying a low-pass filter below the carrier frequencies. However, in practice, the stimulation signal often surpasses the recording system’s capabilities, leading to saturation and the creation of a spurious signal at the Δf frequency. This complicates the differentiation between a genuine neuronal network response and amplifier saturation. Moreover, this phenomenon highlights a drawback of spTI: the carrier frequencies used for stimulation contain the 50Hz frequency, which can potentially contaminate recordings at the Δf frequency, thereby introducing complexities in the analysis of electrophysiological data within this frequency band.

The discovery of envelop frequency modulation of individual neurons represents a significant advancement, as it is the first time such neuronal activation at the Δf frequency has been empirically demonstrated within the context of TI, and in an imaging modality uncorrupted by stimulation artifact. However, the precise temporal nature of this neural response remains an open question. As calcium indicators are limited by slow decay kinetics it was not possible for us to discern the exact number of times a neuron fired during each of these modulated periods. Here, we may be observing bursts of firing where the frequency of the bursts follows the Δf. It is unlikely though with the lower envelope frequency (e.g. 0.1 Hz) that the neurons are limited to firing only once per envelope given the sustained level of calcium within the cells. Additionally, the decay kinetics limited us to examining only low frequency TI as we did not have the temporal precision to accurately capture modulation at higher frequency. Moving to genetically encoded voltage indicators would greatly advance this work, providing a more direct readout of neuronal activity, and superior temporal resolution.

However, spTI allows for better control over the temporal characteristics and duration of the stimulating signal compared to classic TI, improving the temporal precision and control of stimulation. By incorporating PSK modulation, spTI can create pulses, a stimulation waveform highly used during by DBS stimulation using depth leads. spTI can overcome the ambiguous, gradual stimulation onset in regular TI, which depends on the slope of the envelope of the AM signal. To solve this limitation, the introduction of PSK modulation enables the creation of classic bursts of pulses. By controlling the duration of each pulse independent of the stimulation frequency, we achieve customizable bursts, which is an important feature in clinical research and therapy. This enhancement is a significant advancement, as it extends the reach of non-invasive stimulation to deep brain structures, which were previously challenging to target accurately non-invasively.

In this study, we also address a key limitation of regular TI by introducing a multipolar configuration, which enables precise control over the electric field intensity at the target without compromising the spatial profile of stimulation. Unlike conventional TI, the multipolar approach, termed mpTI, utilizes overlapping multiple foci to achieve a cumulative effect, resulting in a substantial increase in the electric field without the need for higher stimulation from individual electrode pairs. This novel configuration minimizes the risks associated with high stimulation levels while preserving the necessary spatial precision for accurate targeting.

Furthermore, we incorporate Fourier components into the multipolar layout, allowing for the replication of classic square biphasic bursts of square pulses, again, commonly used in invasive DBS. This innovation facilitates a seamless transition from invasive to non-invasive stimulation techniques, broadening the potential applications of TI in clinical research.

To evaluate the effectiveness of these proposed modifications, we conducted experiments using a mouse model of epilepsy. We successfully reconstructed the signals at deep brain stimulation targets, specifically the hippocampus, and compared the efficacy of spTI and TI in inducing AD. The results demonstrate that pulse TI, particularly spTI, exhibits significantly higher efficacy (same effect for ∼30% less applied current) in evoking AD. This finding highlights the potential of spTI as a powerful tool for studying epilepsy and other neurological disorders.

Overall, our study introduces a novel multipolar configuration that enhances the versatility and precision of TI stimulation, where mTI and mpTI seems to be equivalent with regard to their depth profile.In conclusion, this paper presents a compelling argument for the effectiveness of multi-pulse TI as a tool for clinical research. The proposed modifications, including the use of PSK modulation, the implementation of a multipolar configuration, and the replication of classic DBS stimulation patterns, address the limitations of traditional TI and enhance its ability to provide targeted and deep brain stimulation. The results obtained in a mouse demonstrate the potential of multipolar technique to achieve a stronger effect at depth with the same applied current at the electrode surface. Further research and validation in human subjects will be crucial to fully establish the clinical utility of this innovative approach.

## Acknowledgments

A.W. received funding from the European Union’s Horizon Europe research and innovation programme under grant agreement No. 101101040 (TREATMENT) and No. 101088623 (EMUNITI). Funding for this work was also provided in part by the United States National Institute of Health (F31NS115479 [MAS], S10OD021773 [KB]) and the Mirowski Family Foundation (REG).

## Authors contribution

A.W. conceived the project with E.G. and A.G.P.

E.A., B.B., D.D., M.A.S. E.C. F.M. C-A.G. M.G. performed experiments and E.A., K.B, M. A.S. and E.C. performed and analyzed calcium imaging experiments. DD contributed to the conceptualization, design and implementation of the spTI and mpTI paradigms based on the PSK and PM techniques and the arbitrary waveform shape TI based on the Fourier series representation. E.N. M.S and A.C. did the finite element modeling and exposure metric discussion. E.A. and A.W. wrote the paper with input from the other authors including V.J, R.E.G and DLD.

## Ethical Publication Statement

We confirm that we have read the Journal’s position on issues involved in ethical publication and affirm that this report is consistent with those guidelines.

## Figures

**Figure supplementary 1:**
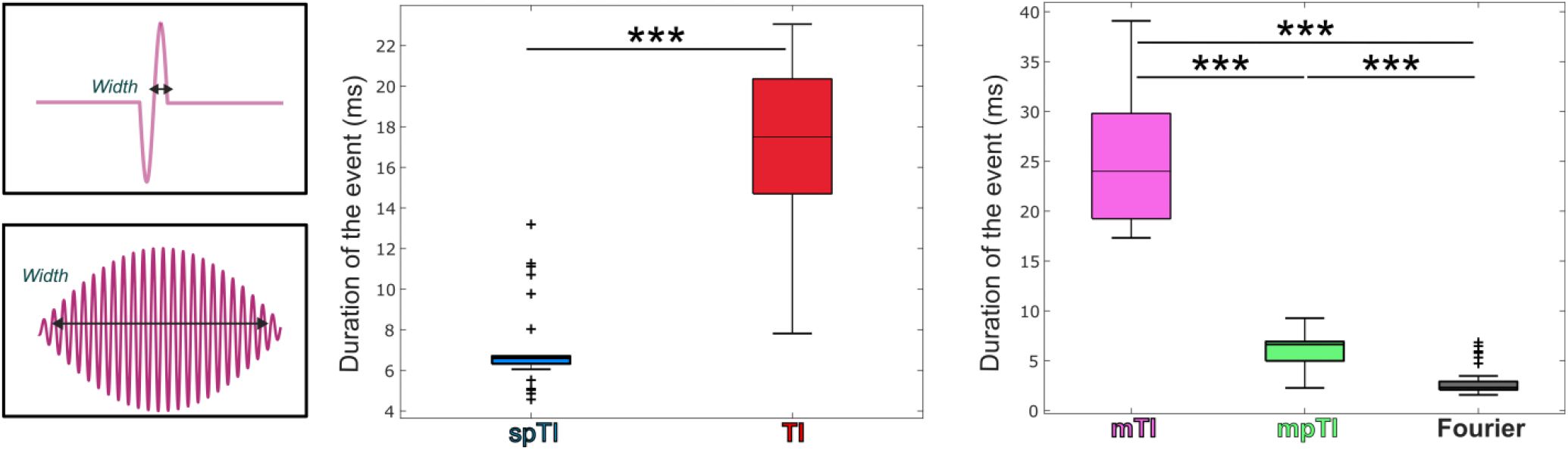
Variation of the duration of the event (either AM or pulses) for each groups. All the groups depending on the AM (TI and mTI) have a significantly longer duration of stimulation event compared to the other groups. p-value*** < 0.001

